# Non-destructive, high-resolution T cell characterization and subtyping via deep-ultraviolet microscopy

**DOI:** 10.1101/2025.05.20.655120

**Authors:** Viswanath Gorti, Caroline E. Serafini, Aaron D. Silva Trenkle, Kaitlyn McCubbins, Isaac LeCompte, Gabriel A. Kwong, Francisco E. Robles

## Abstract

T cell characterization is critical for understanding immune function, monitoring disease progression, and optimizing cell-based therapies. Current technologies to characterize T cells, such as flow cytometry, require fluorescent labeling and typically are destructive endpoint measurements. Non-destructive, label-free imaging methods have been proposed, but face limitations with throughput, specificity, and system complexity. Here we demonstrate deep-ultraviolet (UV) microscopy as a label-free, non-destructive, fast and simple imaging approach for assessing T cell viability, activation state, and subtype with high accuracy. Using static deep-UV images, we characterize T cell viability and activation state, demonstrating excellent agreement with flow cytometry measurements. We further apply dynamic deep-UV imaging to quantify intracellular activity, enabling fast and accurate subtyping of CD4^+^ and CD8^+^ T cells. These results corroborate recent studies on metabolic activity differences between these subtypes, but now with deep-UV microscopy they are enabled by a non-destructive, fast, low-cost and simple approach. Together, our results demonstrate deep-UV microscopy as a powerful tool for high-throughput immune cell characterization, with broad applications in immunology research, immune monitoring, and development of emerging cell-based therapies.

## Introduction

T cells are critical for immune system function, coordinating physiological responses to infections and other immune disorders via cytotoxic activity and cytokine secretion (1, 2). T cells originate as naïve cells, which circulate throughout the host organism and do not express activation markers (e.g., CD25, CD44, and CD69). Upon antigenic exposure, naïve T cells become activated, undergoing rapid proliferation and differentiation into effector T cells (3, 4). These effector cells perform important immune functions such as providing targeted cytotoxicity and helping coordinate immune responses. Over time, some activated T cells differentiate into memory T cells, which remain in the host long-term and enable a rapid response upon subsequent antigen encounters. Monitoring T cell viability, activation state, and subtype - such as CD8^+^ cytotoxic T cells and CD4^+^ helper T cells - is critical for assessing immune health, tracking disease progression, and optimizing therapeutics including adoptive T cell therapies (1, 3, 5, 6). Changes in T cell populations and phenotypes can serve as key biomarkers for conditions including autoimmune disorders, infectious diseases, and cancer. As a result, accurate T cell characterization is important for both clinical diagnostics and development of emerging immunotherapies.

Adoptive T cell therapies, including chimeric antigen receptor (CAR) T cell therapy, have demonstrated clinical success for treating a variety of malignancies (7–9). These therapies rely on ex-vivo culture and expansion of specific T cell populations, requiring characterization of both activation state and subtype to ensure therapeutic efficacy. Notably, recent work has demonstrated that adoptive T cells derived from naïve populations have higher therapeutic potential as compared to T cells that are activated prior to genetic modification (10, 11). Thus, fast, label-free, and non-destructive characterization of T cells is critical to help advance these cell-based therapies.

Current technologies for T cell enumeration and subtyping primarily rely on flow cytometry, a technique that enables high-throughput single-cell analysis. However, there are several limitations to flow cytometry as this technique 1) requires expensive scientific-grade analyzers, 2) uses chemical reagents, and 3) has limited its applicability for real-time monitoring and in-line applications. Recently, label-free optical imaging modalities, including quantitative phase imaging (QPI) and nonlinear optical techniques (e.g., second and third harmonic generation) have been used for *in-vitro* T cell characterization (12, 13). Notably, fluorescence lifetime imaging microscopy (FLIM) has been leveraged to characterize T cells by measuring single-cell metabolic changes via the optical redox ratio (14). This work revealed distinct metabolic signatures between cells of different activation states, demonstrating the potential of optical technologies for robust immune cell characterization. However, these label-free techniques still suffer from constraints in imaging speed, system cost, and complexity, limiting their applicability for high-throughput and accessible T cell characterization.

To address these limitations, we employ deep-UV microscopy, a powerful label-free optical imaging tool that leverages the intrinsic absorption properties of biomolecules in the deep-UV spectrum (200–300 nm) to generate high-contrast images with molecular specificity without any exogenous labels (15). The technique was initially demonstrated for live cell imaging in the early 1900s but has been historically limited by poor detector sensitivity in the deep-UV range and inefficient illumination sources, leading to phototoxicity concerns (16). Recent advancements in UV imaging technology - including improved UV-sensitive detectors, high-efficiency broadband and LED-based illumination sources, and UV-transparent optics - have reinspired interest in the optical technique. In fact, studies have shown that UV exposure typical of deep-UV microscopy studies remains significantly below biological photo-damage thresholds, enabling non-destructive, longitudinal live cell imaging (17). Using quantified absorption spectra of biologically relevant molecules, the technique was successfully applied for quantitative mass mapping of nucleic acids and proteins, which have unique absorption peaks at 255 and 280 nm, respectively (18, 19). More recently, deep-UV microscopy was demonstrated for various hematological applications, including fast, stain-free classification of blood cells using benchtop and compact UV systems, diagnosis of clinical disorders such as neutropenia, adequacy assessment of bone marrow aspirates and virtual histopathology of various tissue types (20– 28).

In this work, we leverage the high-resolution and label-free molecular imaging nature of deep-UV microscopy to image and characterize T cell viability and activation state, demonstrating strong agreement with conventional flow cytometry. Furthermore, we extend the application of UV microscopy by subtyping activated CD4^+^ and CD8^+^ T cells by analyzing dynamic intracellular activity. These measurements reveal functional differences between CD4^+^ and CD8^+^ T cells that correlate with established metabolic differences between the two cell types. Together, this study highlights the potential of deep-UV microscopy as a non-destructive, high-throughput alternative for accurate T cell characterization and subtyping, addressing limitations of existing technologies and advancing immune cell analysis for research and clinical applications.

## Results

### Deep-UV microscopy enables longitudinal imaging of live T cells

To prepare samples for deep-UV microscopy experiments, T cells were longitudinally cultured to cover the T cell lifecycle (Fig. 1A). Naïve CD4^+^ and CD8^+^ T cells were isolated from peripheral blood mononuclear cells (PBMCs) and activated on Day 0 (D0) of culture using CD3/CD28 Dynabeads with IL-2, which mimics the natural activation process.. These cells then transitioned from a naïve state to an activated state by D3, characterized by morphological changes, increased proliferation, and expression of activation markers e.g., CD69 (early activation) and CD25 (sustained activation). Over time, cells transitioned to a post-activated, or contraction phase, state with corresponding morphological changes. These post-activated cells can include phenotypes such as memory T cells; however, this transition can take up to 60 days to occur, and literature has shown that CD8^+^ memory T cell formation requires the presence of CD4^+^ T cells during activation, which was not met in our studies (4, 29, 30). Therefore, for the purpose of this work, we broadly categorize cells post-activation as contraction phase. Flow cytometry was used to monitor activation state during cultures, providing ground-truth validation for deep-UV imaging. CD69 expression was used as an early activation marker for the first few days of culture post-stimulation. CD25 expression was used as a marker of sustained activation, persisting through the activation phase until cells transitioned into contraction phase. By D21, cells exhibited post-activated phenotypes with little to no CD69 or CD25 expression, indicating completion of the activation cycle and progression to contraction phase.

**Fig. 1:**
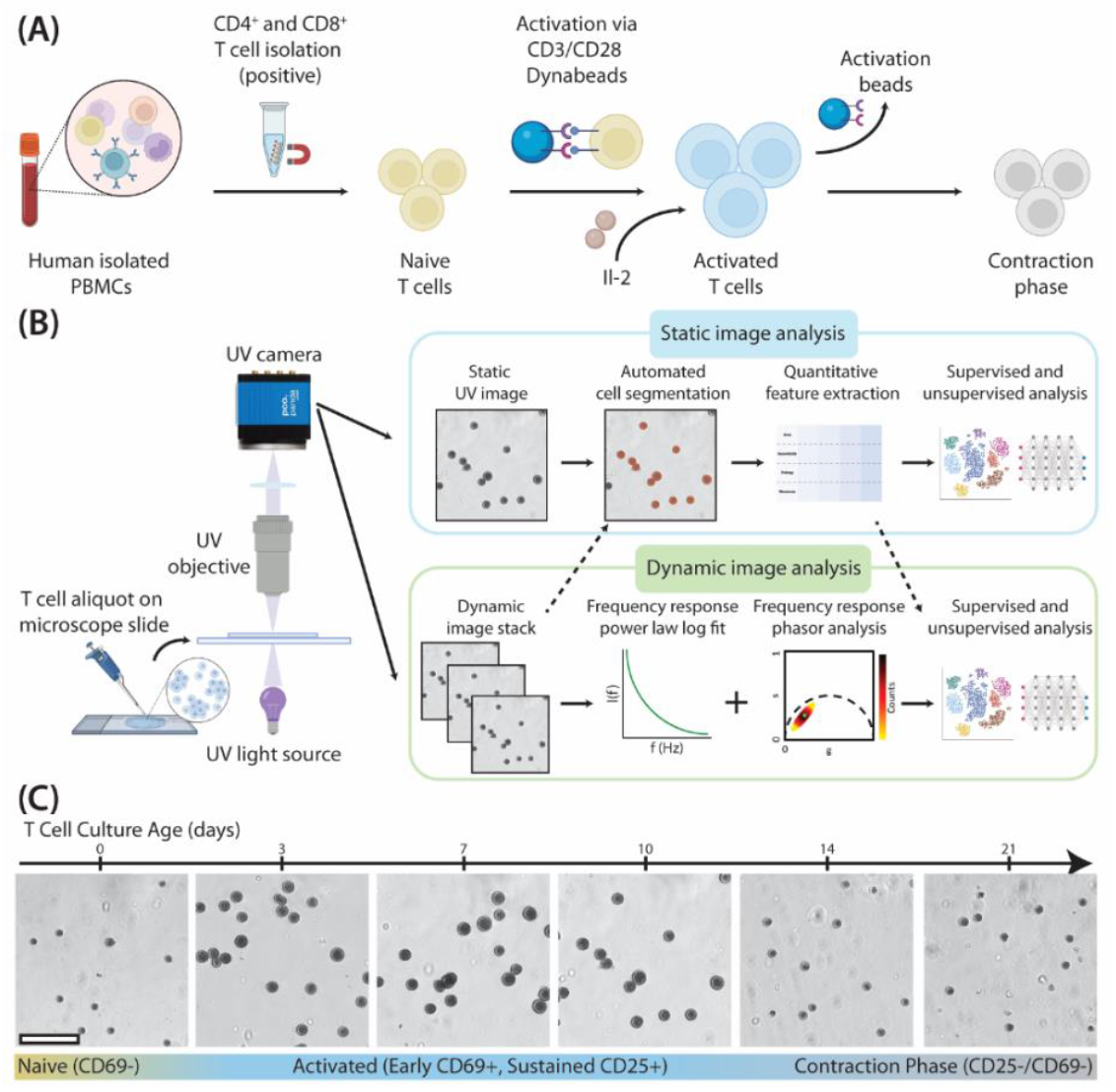
Deep-UV microscopy enables longitudinal imaging of live T cells. (A) Schematic of T cell isolation and activation protocol. (B) Sample preparation, imaging, and analysis workflow using deep-ultraviolet (UV) microscopy. (C) Sample deep-UV images of CD4^+^ T cells during a 21-day culture showing activation and subsequent contraction into post-activated cell phenotypes. Scale bar: 60 µm.

Figure 1B shows the sample preparation, imaging, and analysis workflow for deep-UV microscopy, described in detail in Sections 4.1-4.4 of the Methods Section below. Our approach only requires a small, untreated aliquot of cell suspension to be placed on a microscope slide for imaging at 255 nm illumination (corresponding with the nucleic acid peak in the UV spectrum) (15). This allows for rapid, non-destructive imaging with minimum perturbation to cells, closely preserving their native state. Static (single image) and dynamic (image stack) data of live T cells were captured with a previously demonstrated benchtop deep-UV microscopy setup, comprising a deep-UV light source, UV objective, and UV-sensitive sCMOS camera (20) (Fig. S1A). Static images capture morphological and textural features of cells at a single time point, while dynamic image stacks enable quantification of intracellular activity via frequency response analysis.

Figure 1C shows representative deep-UV images of a sample CD4^+^ expansion taken over a 21-day culture, acquired using a 0.5NA UV objective (resolution of ∼300 nm). As expected, activated T cells exhibit increased cell size and proliferation following activation at D0. This activation persists through D10 of culture (confirmed via flow cytometry), at which point CD25 expression begins to decrease to approximately 0% at D21. This shows that deep-UV microscopy can capture longitudinal morphological changes in live T cell cultures corresponding with activation and post-activation transitions.

### Static deep-UV images enable characterization of T cell viability and activation state

Deep-UV images revealed clear morphological differences between T cell populations, motivating development of automated classification pipelines to characterize T cell viability and activation state. First, a custom residual neural network (detailed in the Methods section) was trained to classify cells into three groups: activated, dead, and quiescent, which comprised both naïve and contraction phase cells. The input data for the network were cropped, segmented, and zero-padded images of segmented T cells from five unique human donors (n = 1200 cells), mixed within an 80/20 train and test dataset split. Ground truth labels for activation state were obtained from flow cytometry. Cells from D0 and D21 were classified as quiescent, with low (<5%) expression of activation markers CD69 and CD25, respectively. Cells from D3-10 were classified as activated, as samples demonstrated >90% CD25 expression. Dead cells were identified using a previously validated, image-based live/dead assay using deep-UV microscopy (17). The viability of each sample assess with the deep-UV images was then compared to flow cytometry. The classification model demonstrated excellent performance, with an overall accuracy of approximately 95% on the test dataset (Fig. 2A).

**Fig. 2:**
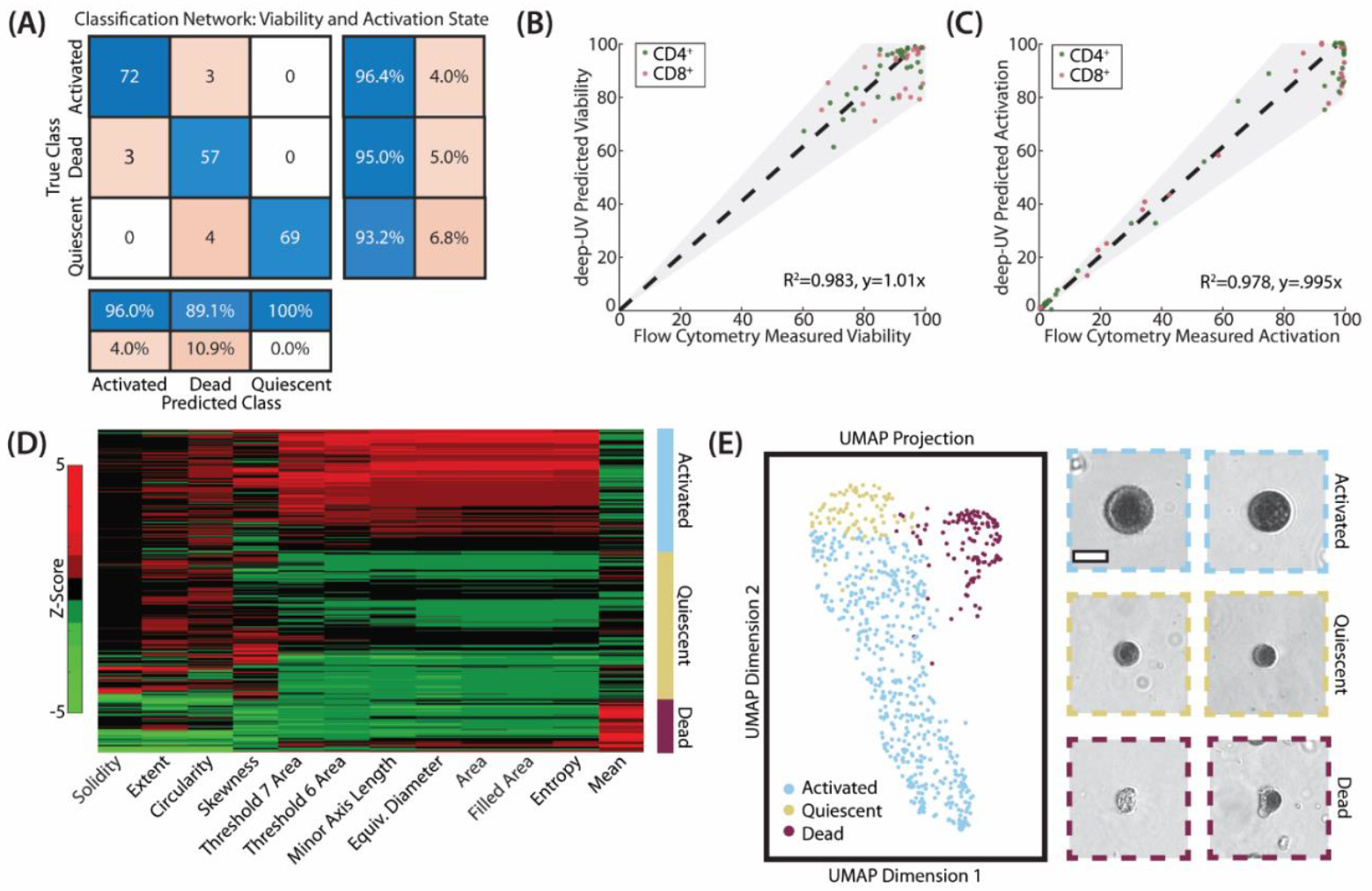
Characterization of T cell viability and activation state via static deep-UV microscopy images. (A) Three-class confusion matrix from a custom residual classification network for characterization of activated, dead, and quiescent cells. Input images were single-channel, segmented UV images of T cells. (B) Scatter plot comparing flow cytometry-derived viability percentage for T cell samples with deep-UV microscopy predicted viability. (C) Scatter plot comparing flow cytometry-derived activation percentage for T cell samples with deep-UV microscopy predicted activation percentage. (D) Clustergram showing the 12 most important quantitative features extracted from static deep-UV images for classification of activated, dead, and quiescent cells. Fractal thresholds 6 and 7 correspond with background corrected image intensity ranges of 0.625-0.750 and 0.750-0.875, respectively. (E) A three-class UMAP showing activated (blue), quiescent (yellow), and dead (red) cells. Sample images of cells from each class are shown on right. N = 1255 cells (5 donors). Scale bar: 10 µm.

To validate its applicability for real-time T cell characterization, we applied the network to previously unseen cells from T cell samples that had been characterized with flow cytometry. These samples comprised T cells isolated from three other human donors. Deep-UV microscopy-predicted viability demonstrated strong agreement with flow cytometry-measured viability for the corresponding samples (Fig. 2B, r^2^ = 0.983, y=1.01x), demonstrating that viability can be assessed from static deep-UV microscopy images. Similarly, deep-UV microscopy-predicted activation percentages for given samples also strongly correlated with flow cytometry-derived activation percentages (Fig. 2C, r^2^ = 0.978, y=0.995x), demonstrating accurate assessment of T cell activation with deep-UV microscopy. We note that classification error may correspond with activation state heterogeneity within T cell cultures, as D3-10 samples did not demonstrate 100% CD69 expression but all cells were assumed to be activated for our study.

To understand differences between T cell phenotypes, we conducted feature-based analysis by extracting a set of morphological and texture-based features from static deep-UV images (detailed in the Methods Section). Feature ranking using chi-squared testing revealed the twelve most relevant features for distinguishing the T cell classes of interest (Fig. S2). Clearly, the mean attenuation of 255 nm light is the best indicator of cell viability, while a combination of cell area, and entropy and fractals—indicative of the intracellular distribution of nucleic acid and to some extent proteins—correlate with T cell activation. The clustergram in Figure 2D displays Z-score distribution across the three groups for each of these features, demonstrating class-specific dependence. Using these features, we leveraged dimensionality reduction via Uniform Manifold Approximation and Projection (UMAP) for visualization of T cell populations. The resulting UMAP showed great separation between activated (blue), quiescent (yellow), and dead (red) cells, demonstrating the significance of the extracted features for T cell characterization. Representative deep-UV images of cells within each class illustrate clear morphological differences between smaller quiescent and larger, highly attenuating activated T cells, as well as features unique to dead cells (e.g., membrane rupture, cytoplasmic leakage, and nuclear condensation). We note that deep-UV microscopy images can be used to further subtype quiescent cells into naïve and contraction phase cells with a different set of features than used for the three-class classification above (Fig. S3), but we elect to simply characterize cells as activated or not for clinically relevant distinction.

### Deep-UV time series reveal differences in dynamic intracellular activity between CD4^+^ and CD8^+^ T cells

Given the ability of static deep-UV images to characterize T cell viability and activation state, we initially attempted to subtype activated CD4^+^ and CD8^+^ T cells using only static images and similar static image features. However, this approach did not yield distinct separation between the two, suggesting that static morphological features alone are insufficient for subtyping classes (Fig. S4A-B). This motivated dynamic deep-UV imaging to capture both morphological and temporally variant intracellular activity within T cell populations.

Here, we acquired 500-image time series of T cells at 255 nm illumination with ∼8 Hz frame rate. Other sets of imaging parameters were tested and are discussed in the Supplemental materials and Discussion (these include different imaging wavelengths, resolution, coherence properties of the light source, and total number of images). Here temporal fluctuations of the attenuation signal, corresponding to intracellular dynamics, were analyzed in the frequency domain (i.e., Fourier transform of the pixel-wise temporal signal, see Fig. 1B), which we observe to exhibit a power-law behavior. Two complementary pixelwise analyses were performed. First, phasor analysis was used, which is a frequency response analysis commonly used in fluorescence lifetime imaging microscopy (FLIM) and other imaging techniques to quantify spectral or temporal fluctuations by decomposing signals into two terms, denoted g and s, which correspond with the real and imaging components of the signal’s Fourier Transform, respectively (31, 32). Phasor g and s values further from the origin (0,0) in phasor space indicating greater temporal fluctuations and thus higher pixelwise activity. The second analysis comprised a fit to a power law decay function, which recent studies using phase contrast have leveraged to extract intracellular dynamic activity correlating with the metabolic behavior of live breast cancer cells (33). By plotting this decay in log-log fashion, a linear region can be extracted whose slope can also serve as a quantitative measure of intracellular dynamic activity. These slope values typically range between one and two, with higher values (more shallow slopes) corresponding with higher measured intracellular activity.

To leverage extracted cellular activity for subtyping, we trained another residual neural network similar to the one used to distinguish between activated and non-activated T cells. However, instead of using inputs comprising single-channel grayscale images containing UV absorption information, we trained the network with four-channel input images, where the channels corresponded with pixelwise UV absorption, phasor g, phasor s, and fitted power law log slope values. These cells were from the same five human donors used for static classification above (n = 625 cells). The network achieved an accuracy of approximately 90% from a five-fold cross-validation (used to mitigate dataset size limitations), which demonstrates the potential of deep-UV microscopy for T cell subtyping (Fig. 3A).

**Fig. 3:**
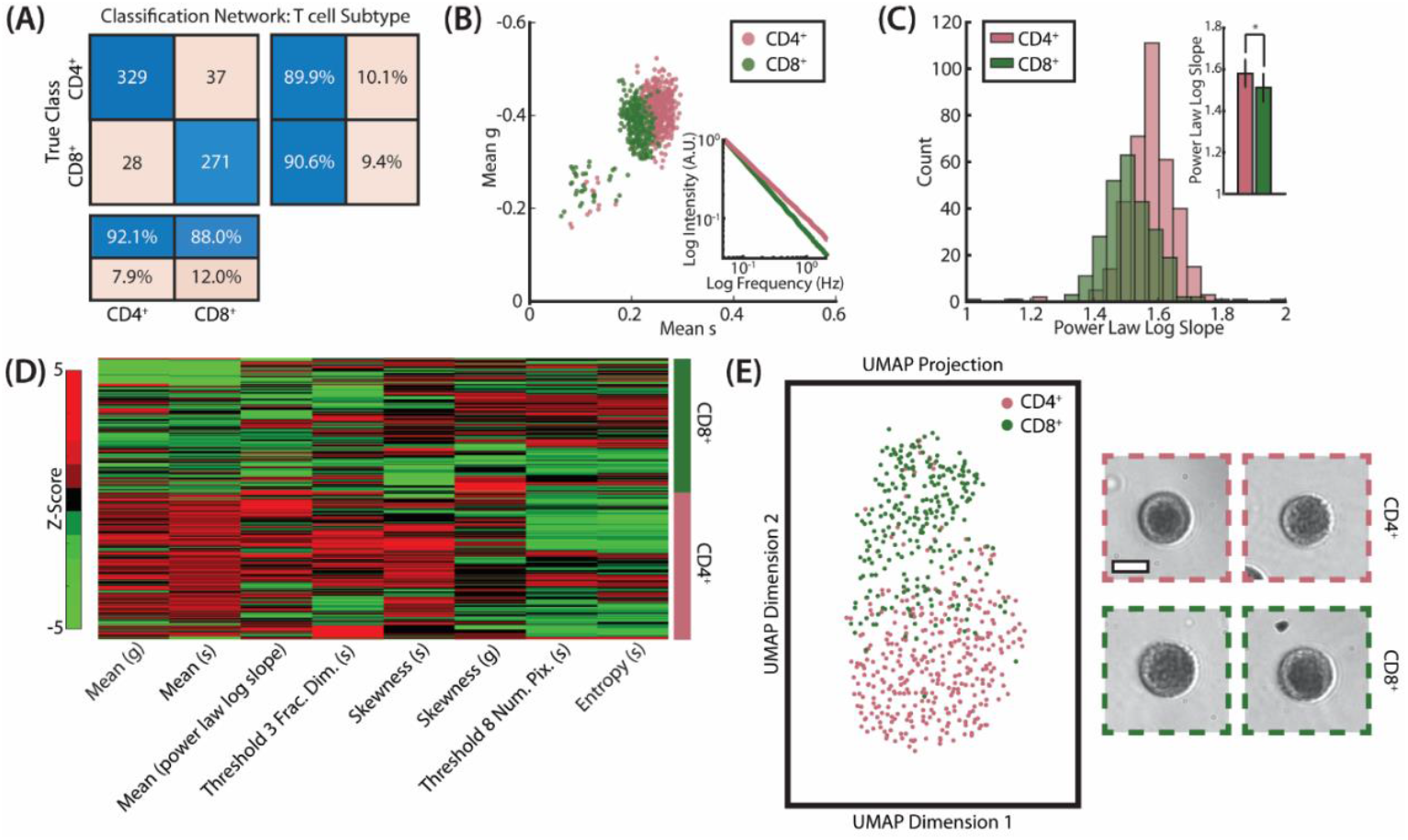
Dynamic frequency response analysis of deep-UV T cell image stacks enables subtyping of CD4^+^ and CD8^+^ T cells. (A) Two-class confusion matrix from a custom residual classification network for classification of activated CD4^+^ and CD8^+^ T cells. Input images were four-channel images containing deep-UV absorption, phasor g, phasor s, and fitted power law log slope images, respectively. (B) Scatter plot of cellwise mean g and s values from phasor analysis for activated CD4^+^ and CD8^+^ T cells. Inset shows a log-log plot of the average dynamic frequency response for each subtype. (C) Histogram of cellwise fitted power law log slope for activated CD4^+^ and CD8^+^ T cells. Inset shows a bar plot of mean slope value for each subtype. * Student’s *t* test p-value <0.05 (D) Clustergram showing the eight most important quantitative features extracted from dynamic deep-UV image stacks for classification of activated CD4^+^ and CD8^+^ T cells. Fractal thresholds 3 and 8 correspond with background corrected image intensity ranges of 0.25-0.375 and 0.875-1, respectively. (E) A two-class UMAP showing CD4^+^ (red) and CD8^+^ (green) cells. Sample images from each class are shown on the right. N = 625 cells (5 human donors). Scale bar: 10 µm.

To understand the source for T cell subtyping, we analyzed cellwise data from our frequency response analyses. A scatter plot of cellwise mean g and mean s values (Fig. 3B) revealed distinct separation between the two subtypes, indicating that differences in measured dynamic using deep-UV microscopy contribute to classification accuracy. The inset in Fig. 3B shows a log-log plot of the mean frequency response of each subtype, also showing separation between the two classes. Notably, CD4^+^ T cells map further from the origin in the phasor scatter plot and have a more shallow (higher) log frequency response slope, indicating higher dynamic activity as compared to CD8^+^ T cells. This separation can be observed in an accumulated pixelwise phasor plot for both subtypes as well (Fig. S4C). Similarly, histograms of cellwise average power law log slope values (Fig. 3C) showed a significant difference in the quantified activity between activated CD4^+^ and CD8^+^ T cells, supporting the finding of subtype-specific intracellular activity. As described above, the power law log slope values provide a quantitative value for dynamic activity; here, CD4^+^ T cells exhibit higher slope values on average, also indicating higher measured dynamic activity (33). A similar feature-based analysis was performed to the three-class classification described above, but with additional dynamic morphological and textural features calculated from phasor g and s images. A clustergram of the top eight features is shown in Figure 3D, and a corresponding two-class UMAP demonstrates excellent separation between activated CD4^+^ and CD8^+^ T cells, as expected. The most important features for subtyping were cell-wise quantifications of measured dynamic activity (e.g., mean phasor g/s values and mean power law log slope), textural features in dynamic cell images (e.g., fractal threshold values), and statistical features of the pixelwise dynamic values (e.g., skewness and entropy). After demonstrating that CD4^+^ and CD8^+^ T cells can be subtyped with deep-UV microscopy, we investigated if these differences were spatially localized within cells. First, we pseudo-colorized deep-UV images of the T cells corresponding with both of our dynamic analyses, with blue and red representing low and high activity, respectively (Fig. 4A). For the fitted power law log slope values, this range spanned 1 to 2, where higher values represent greater pixelwise fluctuations. For phasor analysis, we visualized activity using the product of phasor g and s terms as a scalar for signal variation. Smaller values (closer to the origin in phasor space) indicate low activity, and higher values (further from the origin) indicate high intracellular dynamics (Fig. S4C). These values also correspond with the lower left and upper right of the scatter plot in Fig. 3B, respectively. The pseudo-colorized images reveal that intracellular activity is not homogenously distributed within cells but rather localized to the cell cytoplasm with cell nuclei exhibiting less dynamic activity. To quantify this, we plotted averaged line profiles of the fitted power law log slope across the center of four cells, demonstrating lower measured activity in the nucleus (Fig. 4B).

**Fig. 4:**
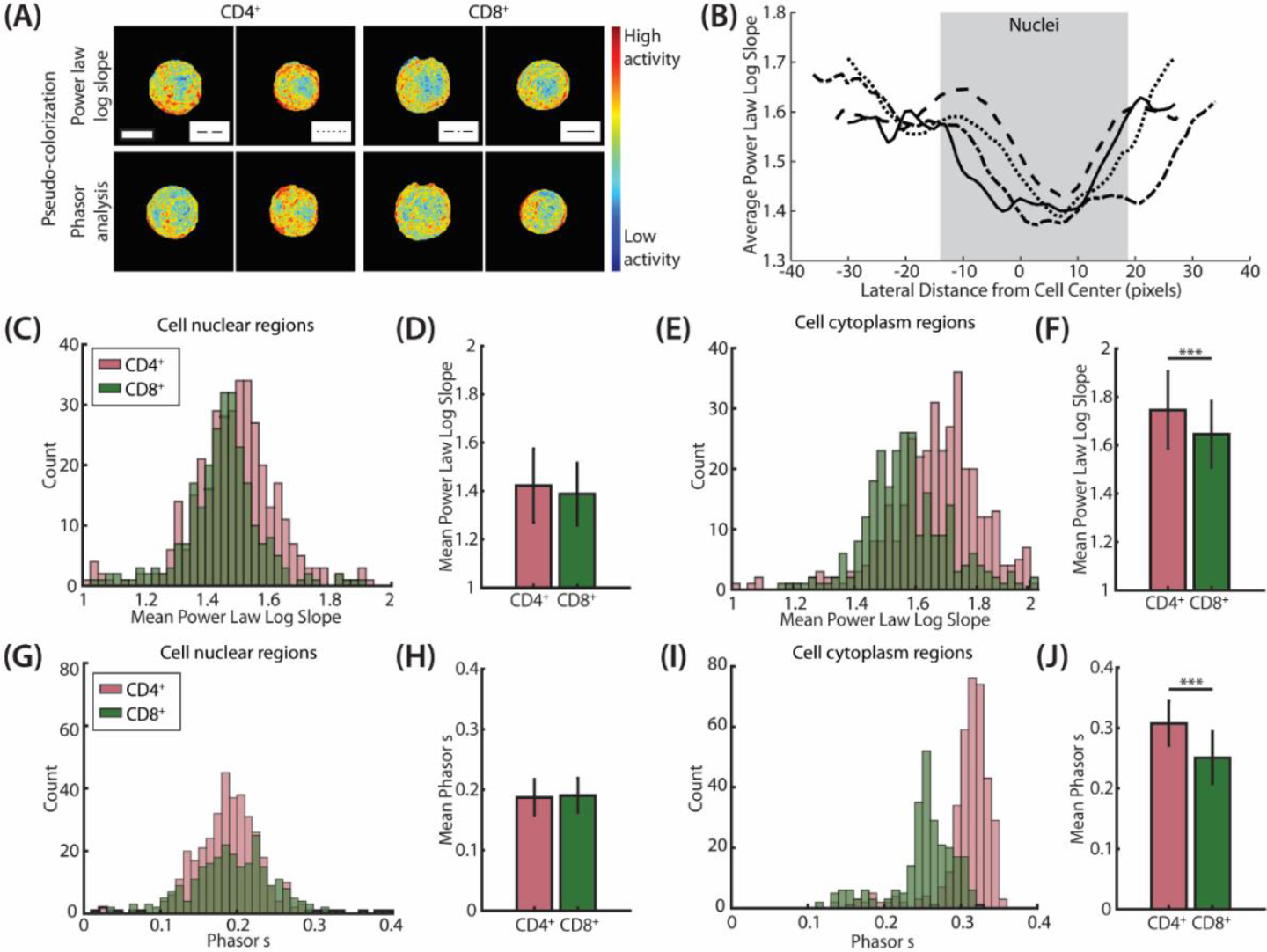
Dynamic deep-UV microscopy image stacks reveal differences in intracellular activity between activated CD4^+^ and CD8^+^ T cells. (A) Sample images of activated CD4^+^ (left) and CD8^+^ (right) T cells pseudo-colorized via fitted power law log slope value (top) and phasor analysis (bottom). Scale bar: 10 µm. (B) Line plots of pixelwise fitted power law log slope averaged over the 10 central horizontal lines of each cell in (A). (C,E) Histograms of fitted power law log slope value for only cell nuclear regions (C) and cell cytoplasm regions (E). (D,F) Corresponding bar plots of mean values of histograms in (C) and (E),*** Student’s *t* test p-value <0.001. (G,I) Histograms of the product of g and s values from phasor analysis for only cell nuclear regions (G) and cell cytoplasm regions (I). (H,J) Corresponding bar plots of mean values of histograms in (G) and (I),*** Student’s *t* test p-value <0.001. N = 625 cells (5 human donors). Scale bar: 10 µm.

To validate this spatial difference in intracellular activity, histograms of the fitted power law log slope were plotted for a set of activated CD4^+^ and CD8^+^ T cells corresponding with segmented cell nuclear regions (Fig. 4C) and cytoplasm regions (Fig. 4E). Nuclear segmentation was performed via image binarization of segmented cells, described further in the Methods Section. These plots demonstrate similar dynamic signatures in cell nuclei between the subtypes, but a significant difference in activity within the cytoplasm, supporting cytoplasmic localization of measured activity. Notably, CD4^+^ T cells demonstrate a higher amplitude of intracellular activity as compared to CD8^+^ T cells, which was also evident when assessing whole-cell dynamics as described above (Fig. 3C). A similar trend is observed with dynamics extracted from phasor analysis, quantified using the cellwise phasor s values (Fig. 4G-J).

## Discussion

In this work, we demonstrated deep-UV microscopy for label-free characterization and subtyping of CD4^+^ and CD8^+^ T cells. By leveraging unique absorption properties of biomolecules in the deep-UV spectrum, we performed high-resolution, non-destructive imaging to classify T cell viability and activation state. Furthermore, we analyzed dynamic intracellular activity to subtype activated T cells into CD4^+^ and CD8^+^ phenotypes with high accuracy, which has not been previously demonstrated with simple, label-free optical systems.

The phenotyping results presented here align with recent studies demonstrating higher metabolic activity in CD4^+^ T cells compared to CD8^+^ T cells. Prior work has shown that CD4^+^ and CD8^+^ T cells have differing metabolic pathways, specifically with respect to reliance on glycolysis and oxidative phosphorylation (34–36). Notably, CD4^+^ T cells exhibit increased glycolysis, oxidative activity, and higher counts of mitochondria within the cytoplasm as compared to CD8^+^ T cells, demonstrated with metabolic assays measuring parameters such as oxygen consumption rate (OCR) and extracellular acidification rate (EAR) (37, 38). Furthermore, studies have found that CD4^+^ T cells demonstrate higher expression of glycolytic enzymes (e.g., hexokinase isoforms) and metabolic complexes (35). This greater reliance on metabolic processes can result in increased organelle motility, intracellular trafficking, and mitochondria remodeling, and has been supported by recent studies investigating immune cell metabolic and mitochondrial dynamics (36–41). Moreover, studies with FLIM have also revealed metabolic differences between the two subtypes via optical redox ratio measurements (14). Given that deep-UV microscopy generates contrast from intrinsic biomolecular absorption, our results suggest that the observed subtype-specific differences in intracellular dynamics correlate with these metabolic differences between cells. Fundamentally, we show that analysis of live cell temporal dynamics enables subtyping of CD4^+^ and CD8^+^ T cells and provides insights into metabolic differences between these phenotypes.

The experiments presented here were conducted using a benchtop deep-UV microscopy system, but similar characterization can be achieved with a previously demonstrated compact, low-cost, portable, and LED-based UV system (Fig. S5). This would enable real-time monitoring of T cell cultures within an incubator or at the point-of-care for applications such as immunotherapy monitoring and personalized medicine. Additionally, we explored the minimum number of dynamic images required for accurate subtyping. While the work shown here used 500 images per time series (similar to previous dynamic imaging studies (42–44)), our results suggest that as few as 300 images – translating to approximately 30 seconds of imaging – may be sufficient for reliable classification and subtyping of CD4^+^ and CD8^+^ T cells (Fig. S6). Moreover, integration of other dynamic analysis pipelines, such as an autocorrelation-based approach that leverages correlation decay analysis to extract dynamic information from fewer time points, could further reduce the number of frames required (45, 46). Together, this would improve classification efficiency and expand the feasibility of deploying deep-UV microscopy for real-time monitoring applications.

To identify the source of our ability to subtype T cells, we investigated several optical parameters, including wavelength, coherence of the illumination source and lateral resolution. For instance, CD4^+^ and CD8^+^ subtyping using 280 nm or 300 nm illumination instead of 255 nm was not successful, indicating that the molecular specificity plays a key role in enabling subtyping. Our investigation also showed that coherence properties of the light source also play an important role. Specifically, spatially incoherent illumination (e.g., via a deep-UV LED and optical diffuser), significantly reduced subtyping contrast as compared to using a more coherent light source (e.g., laser-driven plasma light source or single element LED located a sufficient distance away from the sample) (Fig. S7). This is likely due to the reduced cross-sectional capabilities of coherent illumination, which in this case enable more information about subcellular dynamics to be captured across the cell. These results suggest that some degree of coherence is necessary to generate contrast sufficient for CD4^+^ and CD8^+^ T cell subtyping, and could be extended to other label-free, high-throughput imaging techniques that have been demonstrated for immune cell characterization techniques (47).

Furthermore, we find that lateral resolution is a significant factor for subtyping T cells using measured dynamic intracellular activity. The image stacks used in this work have a diffraction-limited lateral resolution of approximately 300 nm and provide rich subcellular information for identifying CD4^+^ and CD8^+^ phenotypes. However, downsampling these images stacks by a factor of two (corresponding with a lateral resolution of approximately 600 nm), significantly reduces subtyping accuracy (Fig. S7). This is likely because phenotyping is enabled by capturing subcellular behavior from organelle (e.g., mitochondria) motion, and nucleic acid and protein trafficking.

One of the primary limitations of this study is the potential for motion artifacts. Longitudinal time series can be influenced by any lateral or vertical cell drift, limiting subsequent dynamic feature extraction and analysis. In this work, we ensure spatially stationary cells by placing a cell suspension aliquot into a microscope well and allowing cells to settle (for approximately one minute) prior to imaging. This may be more challenging in an in-line flow system but is feasible and has been demonstrated by leveraging microfluidic devices (47).

Furthermore, the scalability and throughput of this approach are limited by dependence on cell density in suspension, which can influence dataset sizes to ensure that a statistically sufficient number of cells are imaged for analysis. Here, we image cells at typical culture concentration (approximately 1x10^6^ cells per mL), without any centrifugation to concentrate cells prior to imaging. However, this can vary with other T cell cultures (e.g., immunotherapy expansions) thus affecting total cell counts during an imaging session. Further optimizing acquisition speeds and developing automated sample preparation pipelines could help improve throughput.

In conclusion, this work demonstrates deep-UV microscopy for label-free, non-destructive characterization and subtyping of CD4^+^ and CD8^+^ T cells. This work has significant implications for optimizing cell-based therapies, improving our ability to monitor disease progression, and improving the basic understanding of immune function. This technology also has broader potential for investigating other immune cell populations, such as dendritic cells, macrophages, and additional T cell phenotypes including exhausted T-cells – a functional state that presents after prolonged response to an antigen without significant morphological changes. Identifying T cell exhaustion, which could be enabled by measurement of dynamic intracellular activity with deep-UV microscopy, would be critical for advancing adoptive T cell therapies during which cells lose their cytotoxic potential despite maintaining the engineered morphology. Furthermore, these studies could provide novel insights into immune cell morphology and activation dynamics under different physiological and pathological conditions, while helping advance the development of other immunotherapies for cancers and autoimmune disorders. As a high-resolution, non-destructive imaging technique, deep-UV microscopy has the potential to be a powerful research and clinical tool in immunology.

## Materials and Methods

### T cell isolation, culture, and flow cytometry protocol

Peripheral blood mononuclear cells (PBMCs) were isolated from healthy human peripheral blood (as approved by Georgia Tech and Emory University Institutional Review Boards; IRB #H20288) using a density gradient medium (Lymphoprep; STEMCELL Technologies) and PBMC isolation tubes (SepMate; STEMCELL Technologies). Informed written consent was obtained from all participants prior to sample collection. CD4^+^ and CD8^+^ T cells were then isolated using immunomagnetic positive selection kits (EasySep Human T Cell Isolation Kits; STEMCELL Technologies) and cultured in X-vivo medium supplemented with human AB serum, N-acetyl L-cysteine, 2-mercaptoethanol, and human IL-2.

T cells were activated with CD3/CD28 human activation beads (Dynabeads Human T-Activator; ThermoFisher) at a 3:1 bead-to-cell ratio on Day 0. Beads were removed on Day 7. At designated time points corresponding to deep-UV microscopy sessions, flow cytometry was performed to assess cell viability and phenotype. Cells were stained with a dead cell stain (LIVE/DEAD Aqua; ThermoFisher) for viability assessment, and with antibodies against CD3 (BV711, UCHT1 clone; BioLegend), CD4 (FITC, RPA-T4 clone; BioLegend), CD8 (APC, SK1 clone; BioLegend), CD25 (PE, M-A251 clone; BioLegend), and CD69 (BV241, FN50 clone; BioLegend).

### Deep-UV microscopy workflow

For deep-UV microscopy, a 10 µL aliquot of T cell suspension was aspirated from the culture plate onto a microscope slide with a liquid-holding well sticker (Disposable Liquid-Holding Slide Wells; Electron Microscopy Sciences). A coverslip was placed over the sample before imaging in a benchtop deep-UV microscopy system. This custom benchtop deep-UV microscope, extensively described in prior works, consists of a broadband laser-driven plasma light source (EQ-99X; Energetiq), narrowband bandpass filters corresponding with biomolecular absorption peaks in the UV region (Single Bandpass Filters; Chroma), a 40X UV objective (LMU-40X-UVB; Thorlabs), and a UV-sensitive camera (pco.Ultraviolet; Excelitas). A schematic of the microscope setup is provided in the Supplemental Information (Supplemental info: benchtop UV setup schematic).

### Static image analysis

Static image analysis was performed using single-wavelength deep-UV microscopy images acquired at 255 nm. Automatic cell segmentation was performed in MATLAB using a morphological processing pipeline to isolate cells within each FOV. For each cell, a set of quantitative morphological (e.g., area, circularity, and equivalent diameter) and texture-based (e.g., mean, entropy, and fractal-based features) features were calculated. These features have been extensively detailed in previous works for tissue and cell characterization (48–54). Feature ranking was then performed using chi-square tests to identify features relevant for characterization of T cell viability and activation state. Using these features, feature-based analysis was performed by creating a uniform manifold approximation and projection (UMAP) (umap; Python) to serve as a visualization of cell phenotype populations. UMAPs were generated with approximately default values for initial parameters (e.g., number of nearest neighbors = 15 and minimum distance between points = 0.1).

Additionally, image-based classification was performed using a custom residual network, trained with zero-padded images of segmented cells with an 80/20 dataset split and data augmentation via translation and reflection. The model was trained over ∼30 epochs using a cross-entropy loss function, optimizing for characterization of T cell viability and activation state.

### Dynamic image analysis

Dynamic image analysis was performed using deep-UV image stacks acquired at ∼8 Hz over 500 frames with 255 nm illumination. Similar cell segmentation and feature extraction was performed using the first frame of each image stack. To quantify intracellular dynamic activity, pixelwise frequency response analysis was performed with two methods. First, phasor analysis was performed, during which the frequency response is decomposed into two terms (g, s) representing the real and imaginary parts of the signal’s Fourier Transform, respectively. Phasor analysis is well suited for characterizing decay signals and is commonly used for other applications such as quantifying and visualizing fluorescence lifetime signals. Second, each pixelwise frequency response was fitted to a power law decay function, modelling recent work that has demonstrated a power law relationship between frequency and intracellular activity. This decay follows a linear curve when plotted in loglog format, thus allowing the log slope of the fitted power law function to be extracted as another quantitative parameter for intracellular activity. Here, extremely low frequency activated (<0.1Hz) was excluded as it did not follow the linear model. These pixelwise values (phasor g, phasor s, power law log slope) were averaged for over each cell and used as three additional features during feature-based analysis. Furthermore, pseudo-colorized cell images were created using these dynamic values and a RGB colormap, with blue and red representing low and high amplitudes of measured dynamic activity, respectively. Cell nuclei and cytoplasm were segmented using a simple binarization of each cell with a threshold identified via Otsu’s method. Morphological post-processing was performed on each nuclei mask to remove thresholding artifacts and smooth the nuclei borders.

Image based classification for was performed using a similar residual network to the one used for static image analysis, but with four-channel input images (containing pixelwise intensity, phasor g, phasor s, and power law log slope images). This model was similarly trained, optimizing for subtyping CD4^+^ and CD8^+^ phenotypes.

## Acknowledgements

We gratefully acknowledge the following funding sources: National Institutes of Health National Institute of General Medical Sciences: R35GM147437 (F.E.R); Burroughs Welcome Fund grant: CASI BWF 1014540 (F.E.R.); National Science Foundation grant: NSF CBET CAREER 1752011 (F.E.R.); National Institutes of Health National Institute of Biomedical Imaging and Bioengineering grant: R41-EB035057 (F.E.R.); National Institutes of Health National Heart, Lung, and Blood Institute grant: R43-HL167435 (F.E.R.); Georgia Institute of Technology; National Institute of Allergy and Infectious Diseases grant: R01AI171892 (G.A.K.); National Science Foundation Graduate Research Fellowships Program grant: DGE-2039655 (A.D.S.T); National Institutes of Health Cell and Tissue Engineering grant: T32GM145735 (A.D.S.T)

## Author contributions

V.G., C.E.S., A.S.T., and F.E.R. designed the experiments outlined in this work. V.G., A.S.T., I.L., and K.M. prepared samples for imaging. V.G. and K.M. acquired data. V.G. and C.E.S. prepared and performed data analysis. V.G., C.E.S., and F.E.R prepared all figures and the manuscript. All authors reviewed and approved the final version of the manuscript. Fig. 1 created with BioRender.com.

## Data availability

Data underlying the results presented in this paper are not publicly available at this time but may be obtained from the authors upon reasonable request.

## References

1. M. H. Andersen, D. Schrama, P. thor Straten, J. C. Becker, Cytotoxic T Cells. Journal of Investigative Dermatology 126, 32–41 (2006).

2. D. A. A. Vignali, L. W. Collison, C. J. Workman, How regulatory T cells work. Nat Rev Immunol 8, 523–532 (2008).

3. A. D. Waldman, J. M. Fritz, M. J. Lenardo, A guide to cancer immunotherapy: from T cell basic science to clinical practice. Nat Rev Immunol 20, 651–668 (2020).

4. S. M. Kaech, W. Cui, Transcriptional control of effector and memory CD8+ T cell differentiation. Nat Rev Immunol 12, 749–761 (2012).

5. D. L. Barber, et al., Restoring function in exhausted CD8 T cells during chronic viral infection. Nature 439, 682–687 (2006).

6. S. Tyagarajan, T. Spencer, J. Smith, Optimizing CAR-T Cell Manufacturing Processes during Pivotal Clinical Trials. Molecular Therapy - Methods & Clinical Development 16, 136–144 (2020).

7. C. H. June, R. S. O’Connor, O. U. Kawalekar, S. Ghassemi, M. C. Milone, CAR T cell immunotherapy for human cancer. Science 359, 1361–1365 (2018).

8. R. C. Sterner, R. M. Sterner, CAR-T cell therapy: current limitations and potential strategies. Blood Cancer J. 11, 1–11 (2021).

9. D. L. Wagner, et al., Immunogenicity of CAR T cells in cancer therapy. Nat Rev Clin Oncol 18, 379–393 (2021).

10. C. S. Hinrichs, et al., Human effector CD8+ T cells derived from naive rather than memory subsets possess superior traits for adoptive immunotherapy. Blood 117, 808–814 (2011).

11. K. J. Yee Mon, et al., Functionalized nanowires for miRNA-mediated therapeutic programming of naïve T cells. Nat. Nanotechnol. 19, 1190–1202 (2024).

12. D. N. H. Kim, A. A. Lim, M. A. Teitell, Rapid, label-free classification of tumor-reactive T cell killing with quantitative phase microscopy and machine learning. Sci Rep 11, 19448 (2021).

13. E. Gavgiotaki, et al., Detection of the T cell activation state using nonlinear optical microscopy. Journal of Biophotonics 12, e201800277 (2019).

14. A. J. Walsh, et al., Classification of T-cell activation via autofluorescence lifetime imaging. Nat Biomed Eng 5, 77–88 (2021).

15. S. Soltani, A. Ojaghi, F. E. Robles, Deep UV dispersion and absorption spectroscopy of biomolecules. Biomed Opt Express 10, 487–499 (2019).

16. P. E. Hockberger, A history of ultraviolet photobiology for humans, animals and microorganisms. Photochem Photobiol 76, 561–579 (2002).

17. V. Gorti, et al., Quantifying UV-induced photodamage for longitudinal live-cell imaging applications of deep-UV microscopy. Biomed. Opt. Express, BOE 16, 208–221 (2025).

18. B. J. Zeskind, et al., Nucleic acid and protein mass mapping by live-cell deep-ultraviolet microscopy. Nat Methods 4, 567–569 (2007).

19. M. C. Cheung, J. G. Evans, B. McKenna, D. J. Ehrlich, Deep ultraviolet mapping of intracellular protein and nucleic acid in femtograms per pixel. Cytometry A 79, 920–932 (2011).

20. A. Ojaghi, et al., Label-free hematology analysis using deep-ultraviolet microscopy. PNAS 117, 14779–14789 (2020).

21. N. Kaza, A. Ojaghi, F. E. Robles, Hemoglobin quantification in red blood cells via dry mass mapping based on UV absorption. J Biomed Opt 26, 086501 (2021).

22. N. Kaza, A. Ojaghi, F. E. Robles, Virtual Staining, Segmentation, and Classification of Blood Smears for Label-Free Hematology Analysis. BME Frontiers 2022 (2022).

23. A. Ojaghi, et al., Label-free deep-UV microscopy detection and grading of neutropenia using a passive microfluidic device. Opt. Lett. (2022).

24. V. Gorti, N. Kaza, E. K. Williams, W. A. Lam, F. E. Robles, Compact and low-cost deep-ultraviolet microscope system for label-free molecular imaging and point-of-care hematological analysis. Biomed. Opt. Express, BOE 14, 1245–1255 (2023).

25. S. Soltani, et al., Prostate cancer histopathology using label-free multispectral deep-UV microscopy quantifies phenotypes of tumor aggressiveness and enables multiple diagnostic virtual stains. Sci Rep 12, 9329 (2022).

26. S. Soltani, B. Cheng, A. O. Osunkoya, F. E. Robles, Deep UV Microscopy Identifies Prostatic Basal Cells: An Important Biomarker for Prostate Cancer Diagnostics. BME Frontiers 2022 (2022).

27. B. D. Cikaluk, et al., Simultaneous deep ultraviolet transmission and scattering microscopy for virtual histology. Opt Lett 49, 2729–2732 (2024).

28. V. Gorti, et al., Rapid, point-of-care bone marrow aspirate adequacy assessment via deep ultraviolet microscopy. Laboratory Investigation 0 (2025).

29. E. M. Janssen, et al., CD4+ T cells are required for secondary expansion and memory in CD8+ T lymphocytes. Nature 421, 852–856 (2003).

30. B. J. Laidlaw, J. E. Craft, S. M. Kaech, The multifaceted role of CD4+ T cells in CD8+ T cell memory. Nat Rev Immunol 16, 102–111 (2016).

31. M. A. Digman, V. R. Caiolfa, M. Zamai, E. Gratton, The Phasor Approach to Fluorescence Lifetime Imaging Analysis. Biophysical Journal 94, L14–L16 (2008).

32. S. Ranjit, L. Malacrida, D. M. Jameson, E. Gratton, Fit-free analysis of fluorescence lifetime imaging data using the phasor approach. Nat Protoc 13, 1979–2004 (2018).

33. Á. Cano, et al., Rapid mechanical phenotyping of breast cancer cells based on stochastic intracellular fluctuations. iScience 27 (2024).

34. S. Ma, Y. Ming, J. Wu, G. Cui, Cellular metabolism regulates the differentiation and function of T-cell subsets. Cell Mol Immunol 21, 419–435 (2024).

35. Y. Cao, J. C. Rathmell, A. N. Macintyre, Metabolic Reprogramming towards Aerobic Glycolysis Correlates with Greater Proliferative Ability and Resistance to Metabolic Inhibition in CD8 versus CD4 T Cells. PLOS ONE 9, e104104 (2014).

36. C. S. Palmer, et al., Regulators of Glucose Metabolism in CD4+ and CD8+ T Cells. International Reviews of Immunology 35, 477–488 (2016).

37. N. Jones, et al., Metabolic Adaptation of Human CD4+ and CD8+ T-Cells to T-Cell Receptor-Mediated Stimulation. Front. Immunol. 8 (2017).

38. M. Reina-Campos, N. E. Scharping, A. W. Goldrath, CD8+ T cell metabolism in infection and cancer. Nat Rev Immunol 21, 718–738 (2021).

39. C. S. Palmer, M. Ostrowski, B. Balderson, N. Christian, S. M. Crowe, Glucose metabolism regulates T cell activation, differentiation, and functions. Front Immunol 6, 1 (2015).

40. R. Elhage, et al., Mitochondrial dynamics and metabolic regulation control T cell fate in the thymus. Front. Immunol. 14 (2024).

41. E. M. Steinert, K. Vasan, N. S. Chandel, Mitochondrial Metabolism Regulation of T Cell-Mediated Immunity. Annu Rev Immunol 39, 395–416 (2021).

42. J. Scholler, et al., Dynamic full-field optical coherence tomography: 3D live-imaging of retinal organoids. Light Sci Appl 9, 140 (2020).

43. P. C. Costa, et al., Functional imaging with dynamic quantitative oblique back-illumination microscopy. JBO 27, 066502 (2022).

44. C. Filan, et al., Label-Free Functional Analysis of Root-Associated Microbes with Dynamic Quantitative Oblique Back-illumination Microscopy. Res Sq rs.3.rs-3517586 (2023). 10.21203/rs.3.rs-3517586/v1.

45. Dynamic optical coherence tomography algorithm for label-free assessment of swiftness and occupancy of intratissue moving scatterers. Available at: https://arxiv.org/html/2412.09351v1 [Accessed 21 February 2025].

46. C. Ren, et al., Dynamic contrast optical coherence tomography (DyC-OCT) for label-free live cell imaging. Commun Biol 7, 1–8 (2024).

47. C. E. Serafini, et al., Label-Free In-Line Characterization of Immune Cell Culture using Quantitative Phase Imaging. (2025).

48. R. M. Haralick, K. Shanmugam, I. Dinstein, Textural Features for Image Classification. IEEE Transactions on Systems, Man, and Cybernetics SMC-3, 610–621 (1973).

49. An Efficient Algorithm for Fractal Analysis of Textures | IEEE Conference Publication | IEEE Xplore. Available at: https://ieeexplore.ieee.org/document/6382737 [Accessed 21 February 2025].

50. C. T. Jr, A. Traina, L. Wu, C. Faloutsos, Fast feature selection using fractal dimension. Journal of Information and Data Management 1, 3–3 (2010).

51. R. C. Gonzalez, R. E. Woods, Digital image processing, Fourth edition, global edition (Pearson, 2017).

52. F. E. Robles, J. W. Wilson, W. S. Warren, Quantifying melanin spatial distribution using pump-probe microscopy and a 2-D morphological autocorrelation transformation for melanoma diagnosis. J Biomed Opt 18, 120502 (2013).

53. F. E. Robles, et al., Pump-probe imaging of pigmented cutaneous melanoma primary lesions gives insight into metastatic potential. Biomed. Opt. Express, BOE 6, 3631–3645 (2015).

54. P. C. Costa, et al., Towards in-vivo label-free detection of brain tumor margins with epi-illumination tomographic quantitative phase imaging. Biomed Opt Express 12, 1621–1634 (2021).

