## Supplemental Information for "Non-destructive, high-resolution T cell characterization and subtyping via deep-ultraviolet microscopy"

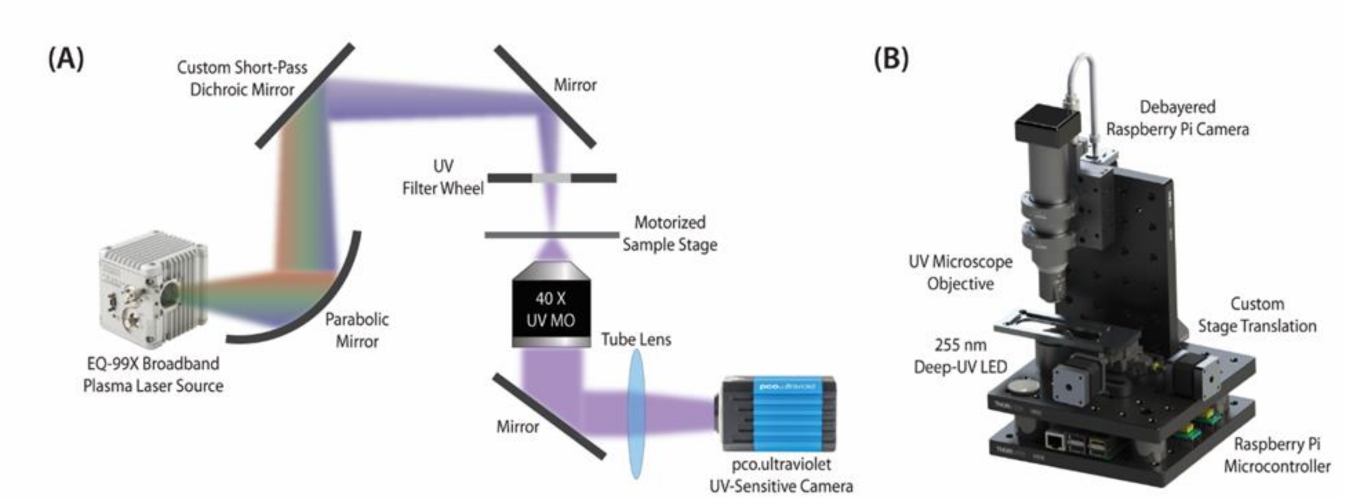


**Fig. S1. Ultraviolet (UV) microscopy setups.** (A) Benchtop plasma laser source-based multispectral UV microscope. Adapted from Ref. (1). (B) Custom compact, deep-UV LED-based microscope system. Adapted from Ref. (2).

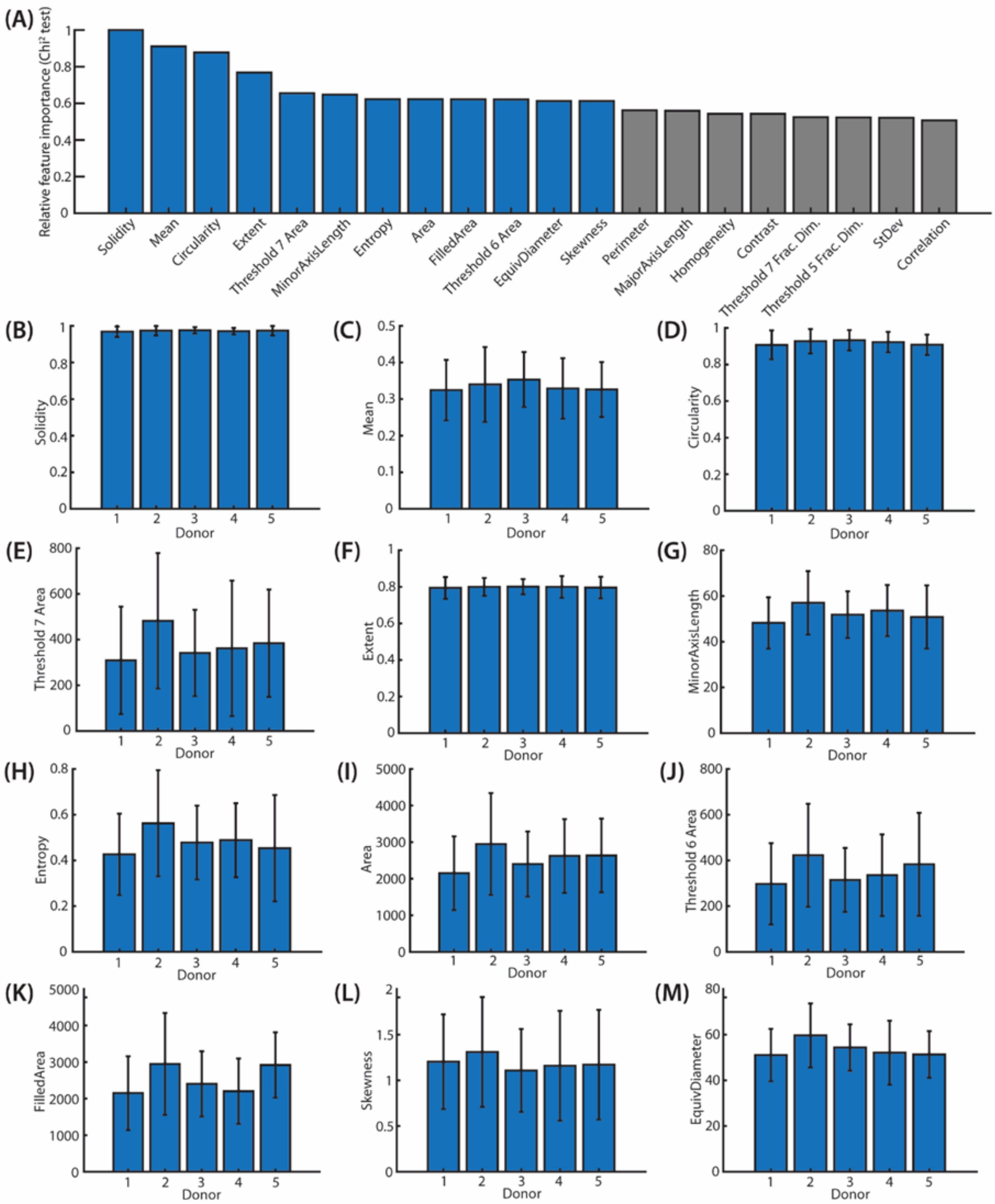


**Fig. S2. Static feature extraction for T cell viability and activation.** (A) Relative feature importance from $\chi^2$testing of selected features extracted from static deep-UV images for classification of T cell viability and activation state. Features with blue bars were used to create the corresponding UMAP. (B-M) Bar plots by donor for the top features used for UMAP visualization. Error bars represent standard deviate

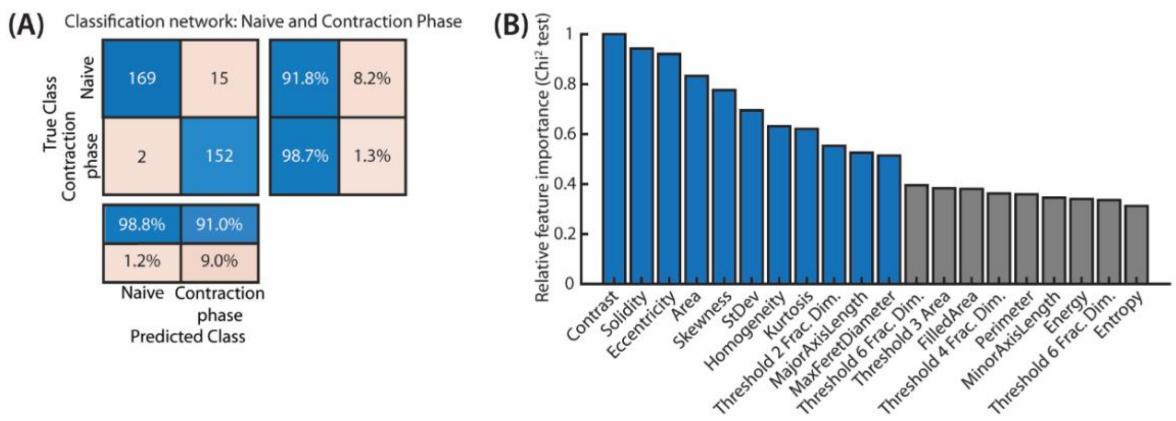


**Fig. S3. Classification of quiescent cells using static deep-UV images.** (A) Confusion matrix from a Residual Network trained for classification of contraction phase and naïve T cells. Dataset comprised 338 single-channel, segmented deep-UV images of quiescent T cells. (B) Relative feature importance from $\chi^2$testing of selected features extracted from static deep-UV images for classification of contraction phase and naïve T cells. Top 11 features indicated with blue bars.

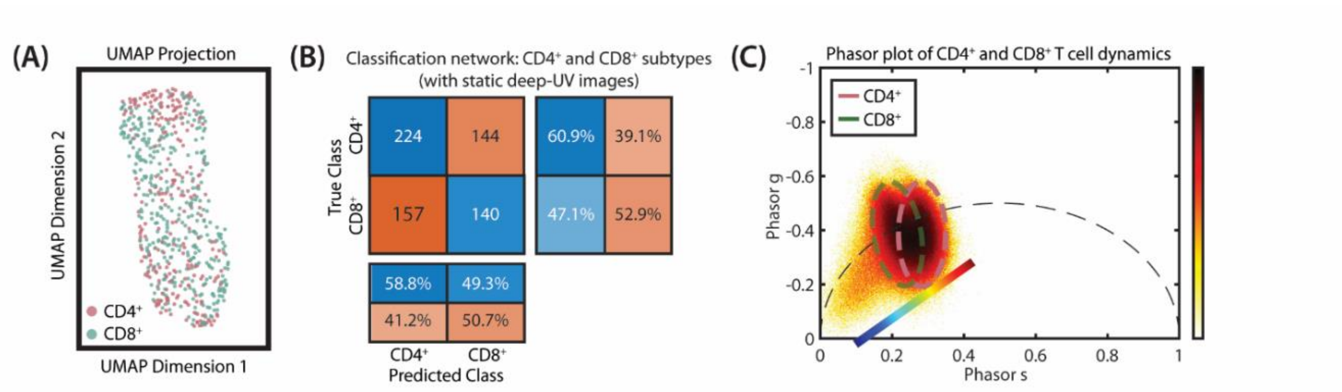


**Fig. S4. Dynamic image data is required for subtyping CD4+ and CD8+ T cells.** (A) A 2-class UMAP of activated CD4+ and CD8+ T cells using the features used for classification of T cell viability and activation state (Fig. S2A). (B) Confusion matrix from a 2-class residual network trained for classification of CD4+ and CD8+ T cells. Training data comprised 605 single channel, segmented deep-UV images of activated T cells. (C) Accumulated pixelwise phasor plot of 605 activated CD4+ and CD8+ T cells, captured with 255 nm illumination and 8 Hz imaging frame rate. Dashed ovals represent the phasor clusters corresponding with CD4+ (pink) and CD8+ (green) T cells. The included diagonal colormap represents the pseudo-colorization scheme used in Fig. 4A.

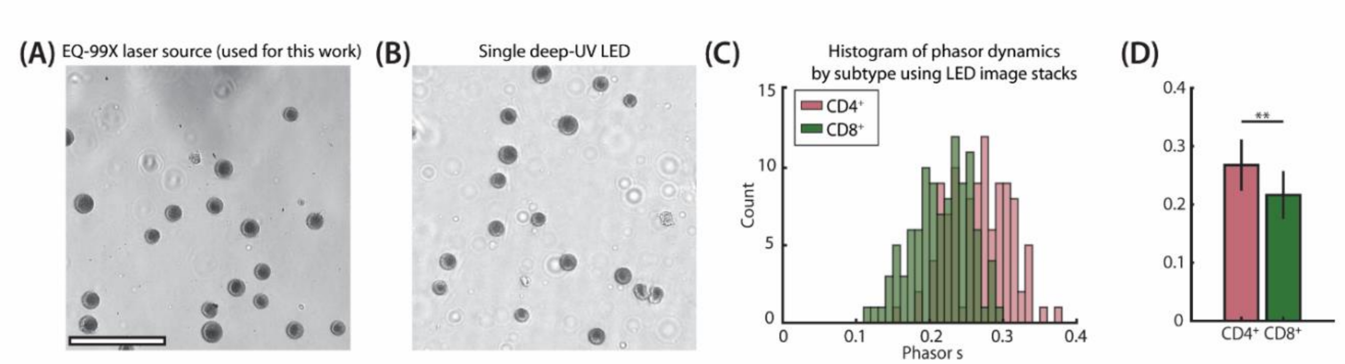


**Fig. S5. LED-based UV microscopy systems enable T cell characterization.** (A-B) Sample images of T cells with 255 nm illumination via the EQ-99X plasma laser-based source used in this study (A) and a single 255 nm deep-UV LED (B). Scale bar: 60 µm. (C) Histogram of cell-wise phasor s values from dynamic image stacks of activated $CD4^+$ and $CD8^+$ T cells. (D) Bar plot of the mean phasor s value for $CD4^+$ and $CD8^+$ T cells from (C). ** Student's *t* test p-value <0.01.

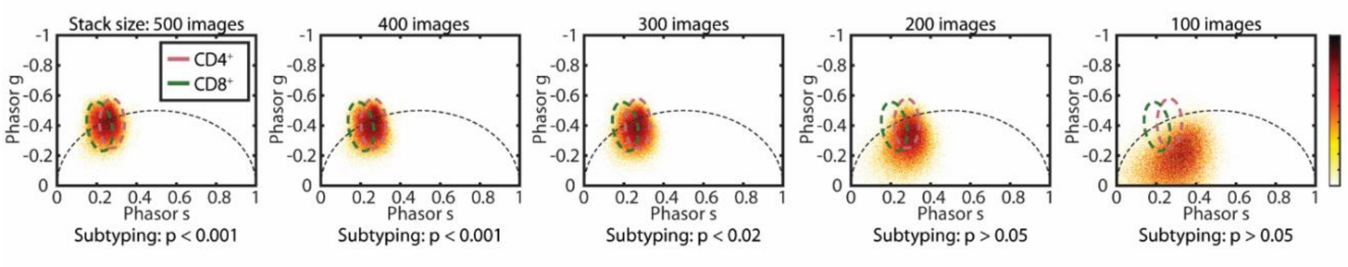


**Fig. S6. Measured intracellular dynamics and ability to subtype CD4+ and CD8+ T cells depend on dynamic image stack size.** Accumulated pixelwise phasor plots shown for activated CD4+ and CD8+ T cells with dynamic image stack sizes of 500, 400, 300, 200, and 100 images (left to right). Dashed ovals on all plots represent the phasor clusters corresponding with CD4+ (pink) and CD8+ (green) T cells. Corresponding p-value from a Student's t test between mean CD4+ and CD8+ cell-wise phasor s values included below each plot.

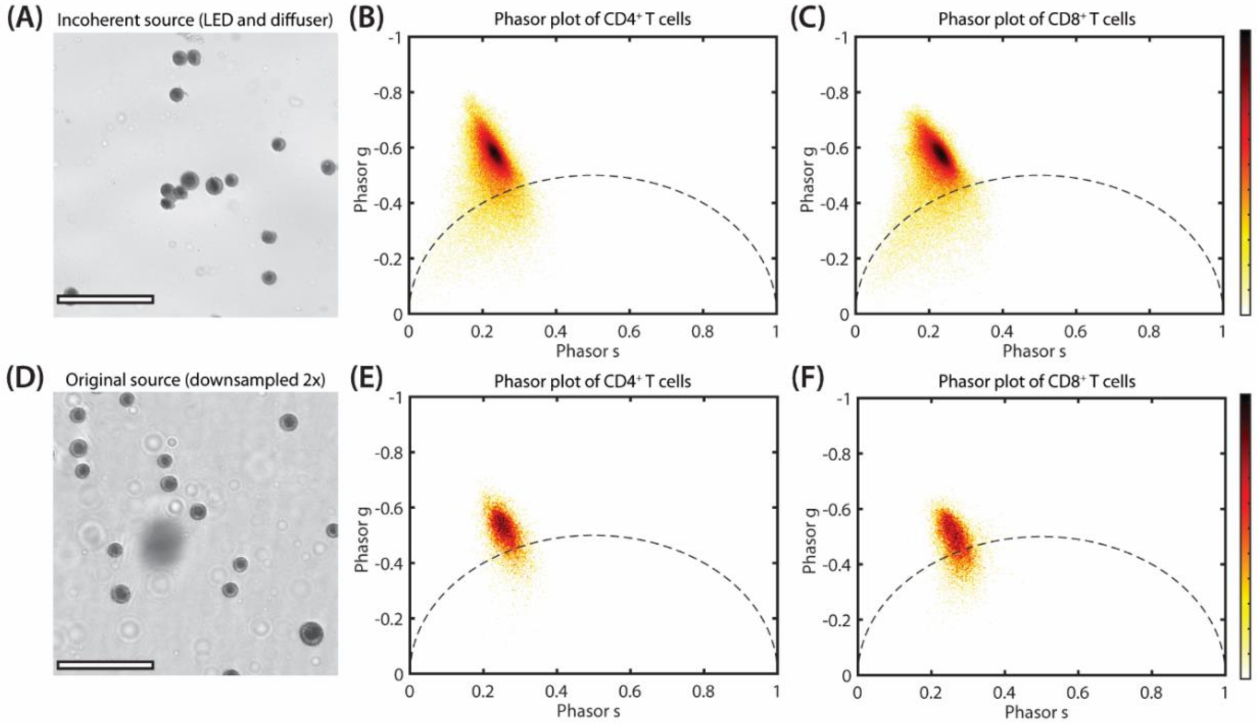


**Fig. S7. Illumination coherence and system lateral resolution affect measured dynamics and subtyping ability.** (A) Sample image of T cells with incoherent 255 nm illumination via a single deep-UV LED and UV fused silica optical diffuser (DGUV10-600; Thorlabs). Scale bar: 60 µm. (B-C) Pixelwise phasor plots of $CD4^+$ (B) and $CD8^+$ (C) T cells from image stacks with incoherent illumination (n = 80 cells per group). Student's *t* test p-value of cell-wise phasor s values >0.05. (D) Sample image of T cells digitally downsampled by a factor of 2, corresponding with a lateral resolution of ~600 nm. Scale bar: 60 µm. (E-F) Pixelwise phasor plots of $CD4^+$ and $CD8^+$ T cells from downsampled image stacks (n = 50 cells per group). Student's *t* test p-value of cell-wise phasor s values >0.05.
